# Sniffing behavior of semi free-ranging Barbary macaques (*Macaca sylvanus*)

**DOI:** 10.1101/2022.08.08.503203

**Authors:** Miriam Simon, Anja Widdig, Brigitte M. Weiß

## Abstract

Olfaction is one of the evolutionarily oldest senses and plays a fundamental role in foraging and social interactions across mammals. In primates, the role of olfaction is now well recognized, but better investigated in strepsirrhine and platyrrhine primates than in catarrhines. We observed the sniffing behavior of semi-free ranging Barbary macaques, *Macaca sylvanus*, at Affenberg Salem, Germany, to assess how frequently macaques sniff and in which contexts, and how sniffing is affected by sex and age. Focal observations of 24 males and 24 females aged 1 to 25 years showed that Barbary macaques sniffed, on average, 5.3 times per hour, with more than 80% of sniffs directed at food. Irrespective of the context, younger individuals sniffed more often than older ones. Females’ sniffs were more often directed at food than male sniffs, while males sniffed more often than females in a social context. Sniffs at conspecifics occurred primarily in a sexual context, with 70% of social sniffs directed at female anogenital swellings performed by males. Of the observed 176 anogenital inspections, 51 involved sniffing of the swelling. Olfactory inspections were followed by copulation significantly less often than merely visual inspections, suggesting that anogenital odors provided additional information guiding male mating decisions. In sum, results show that Barbary macaques routinely use olfaction during feeding, but also in a social context. Our study further suggests that odors may guide mating decisions, but the role of olfaction in sexual interactions warrants further investigations.

## Introduction

Primates, like other mammals, use various sensory modalities to gain information about their social and physical environment. Olfaction is one of the evolutionarily oldest senses and represents an important pathway of information transfer that is used in fundamental behaviors such as foraging (e.g. capuchin monkeys (*Cebus imitator*), Melin et al. 2019), predator avoidance (e.g. mouse lemurs (*Microcebus murinus*), Suendermann et al. 2008) or mating (e.g. ring-tailed lemurs (*Lemur catta*), Boulet et al. 2010).

Among primates, optic convergence and stereoscopic vision represent significant parts of their evolution, accompanied by a relative shrinking of the olfactory apparatus (Kay 2018). Thus, it was long assumed that olfaction is of little relevance in taxa with advanced visual capabilities. Accordingly, the role of olfaction has been studied more intensely in strepsirrhine primates, which are often nocturnal, possess less specialized vision and more developed olfactory structures (Barton 2006) than other taxa. For example, it is well-established that odor plays an important role in their sociality by providing information about sex, age or group membership (Janda et al. 2019; Scordato & Drea 2007) as well as in their foraging behavior (Cunningham et al. 2021; Rushmore et al. 2012). However, it has become increasingly evident that diurnal primates, including haplorrhine primates, may rely strongly on olfaction in a range of contexts. In platyrrhines particularly, several species use active olfaction (i.e. sniffing) in feeding situations, and olfaction may be modulated by visual information. For example, white-faced capuchins (*Cebus capucinus*) and spider monkeys (*Ateles geoffroyi)* sniff fruits that give cryptic visual cues about their ripeness more often than less cryptic ones (Nevo & Heymann 2015; Hiramatsu et al. 2009). Moreover, olfaction also plays a role in the sociality of platyrrhine species, including in the context of mating. For instance, anogenital odor secretions of female common marmosets (*Callithrix jacchus*) vary with fertile phases and males inspect peri-ovulatory odors more intensely than odor samples from follicular and luteal phases (Kücklich et al. 2019).

Catarrhine primates show trichromatic vision in both sexes along with only half the number of olfactory receptor genes and relatively smaller olfactory bulbs compared to strepsirrhine primates (Niimura et al. 2018). In this taxonomic group, the assumption of ‘microsmatic’ primates has persisted over decades (Smith & Bhatnagar 2004). However, a more recent viewpoint suggests that features like the number of olfactory genes are not always predictive of olfactory sensitivity (Smith & Bhatnagar 2004; Laska et al. 2007a; Matsui et al. 2010) and that olfactory behavior is prevalent in many species (Jänig et al. 2018; Vaglio et al. 2021). Furthermore, the importance of proportionally smaller olfactory bulbs has been questioned, because the mere proportional correlation of body size to organ size and, therefore, function may not apply in olfactory systems (Smith & Bhatnagar 2004). In addition, the traditionally assumed separation of the main and the accessory olfactory system appears to be vaguer than previously thought (Sipos et al. 1995; Petrulis et al. 1999).

Hence, it is not surprising that research has also started to focus on the involvement of olfaction in catarrhine sociality and ecology (Matsumoto-Oda et al. 2007; Setchell et al. 2010; Vaglio et al. 2021; Rigaill et al. 2022). For instance, rhesus macaques (*Macaca mulatta*) as well as chimpanzees (*Pan troglodytes*) were shown to differentiate between group and non-group members when presented with samples of body odor (Henkel et al. 2015; Henkel & Setchell 2018). In mandrills (*Mandrillus sphinx*), group members infected with parasites were discovered to have a different fecal odour than non-infected members, whereby healthy individuals avoided the fecal matter of infected conspecifics and avoided grooming them (Poirotte et al. 2017). Both in the wild and in captivity, chimpanzees were observed to sniff at food as well as at conspecifics. While females used olfaction to inspect food more frequently than males did, males were observed to sniff more often in a social and sexual context than females, which was attributed to their fission-fusion society and the high level of male competition (Matsumoto-Oda et al. 2007; Jänig et al. 2018). Hence, evidence is accumulating that catarrhine primates also rely on olfaction in a range of contexts, but the number of studies and species investigated remains low, thereby hampering a more general understanding of the role of olfaction in catarrhine lives.

To contribute to closing the knowledge gap regarding olfaction in catarrhine species, the present study investigated the natural sniffing behavior of Barbary macaques (*Macaca sylvanus*), a catarrhine species native to northern Africa and Southern Europe. Their diet consists of various plants as well as insects (Fooden 2007). Like many other catarrhine species, Barbary macaques live in multi-male, multi-female groups and show a promiscuous mating system (Modolo & Martin 2008). They are sexually active year-round, but show increased sexual behavior during the mating season from late autumn to late winter (Kuester & Paul 1992). Females show a conspicuous visual fertility signal, the anogenital swelling, which is directly linked to estrogen levels and reaches its maximum size during the most fertile phase (Brauch et al. 2007). Nonetheless, observations of genital inspections by looking, sniffing and touching, as well as mating behavior suggest that males may have more than visual information available to determine female fertility (Young et al. 2013).

Hence, we aimed to investigate sniffing behavior in Barbary macaques in different contexts (feeding, social and non-social), and to examine possible variation in sniffing behavior related to sex and age. We hypothesized that individuals who benefit more from olfactory information than others would sniff more frequently, with distinct difference between sexes, age and context. Similar to a study in chimpanzees (Jänig et al. 2018), we, hence, predicted that 1) male Barbary macaques sniff more in the social context than females, with males sniffing particularly females, given that the anogenital swelling was suggested to be a probabilistic cue of fertility (Brauch et al. 2007) and olfactory information would be beneficial in male mating competition. By contrast, we predicted 2) females to sniff edible items more frequently than males, as females are energetically more constrained, similar to findings in female chimpanzees (Matsumoto-Oda et al. 2007). More stringent food inspection has been shown to benefit the long-term reproductive success of females by lowering contamination risks (Rolff 2002; Poirotte et al., 2019). With regard to age, we predicted 3) younger Barbary macaques to sniff more frequently than older individuals irrespective of the context. Infants and juveniles are in a process of learning about their environment and conspecifics surrounding them. Hence, olfactory information could be beneficial for exploring and gaining experience in all contexts, as observed in great apes (Jänig et al. 2018). We also assessed the role of olfaction in inspection behavior of the female sexual swelling and hypothesized an involvement of olfactory information in male mating decision, as male Barbary macaques are suggested to rely not only on swelling size but other information to recognize the female fertile phase (Young et al. 2013). In particular, we predicted that sniffing in the sexual context is influenced by 4) male age, with olfactory inspections being more prevalent in younger males which may lack the experience in visually assessing anogenital swellings and may therefore be more prone to attend to additional cues. Furthermore, the fertility and reproductive success of female Barbary macaques as well as other primates varies across age, with higher variability in interbirth intervals in young females and a decline in reproductive performance from mid age to old age (Paul et al. 1993, Campos et al. 2022). Accordingly, we also predicted that sniffing in the sexual context is affected by 5) female age, with olfactory inspections being aimed more often towards young and old females whose fertility status may be uncertain to the respective male compared to those of prime age. Finally, olfactory inspections were frequently followed by copulations in olive baboons (*Papio anubis*) when females were in their fertile phase but not when they were post-fertile (Rigaill et al. 2013). Hence, we expected that 6) olfactory inspections would affect copulatory behavior in Barbary macaques, although we were not able to address this aspect in as much detail as by Rigaill et al (2013) given that we did not have systematic data on the fertile state of the females during our study period.

## Methods

### Study site

The study was conducted at Affenberg Salem close to Lake Constance, Germany, which is home to ∼ 200 Barbary macaques living in 20 hectares of fenced forest year-round under near-natural conditions, allowing continuous full contact between animals. The park is open to visitors from March to November. Visitors are restricted to a path in one third of the enclosure, while the monkeys can roam freely across the entire area. For more details on the park, see de Turckheim & Merz (1984). The monkeys feed on natural vegetation and insects and receive daily supplements of fruits and vegetables. Furthermore, wheat is distributed widely on and around the clearings and meadows where the food is distributed. Water is accessible at several ponds and water troughs *ad libitum*. The park is home to three naturally formed social groups, each consisting of 50-70 individuals of both sexes and all age classes. All monkeys are individually identifiable by tattoos and natural markings. To control population size, about two thirds of the adult females receive subcutaneous hormonal implants (Implanon NXT), which contain etonogestrel to inhibit the release of egg cells from the ovaries by reducing the luteinising hormone level and impede entry of sperm into the uterus. Implanted females show cyclical changes in anogenital swelling size similar to non-contracepted females (B.M.W., personal observation). This study was purely observational and in accordance with the legal requirements of Germany, all national and institutional guidelines for the care and use of animals, and adhered to the American Society of Primatologists (ASP) Principles for the Ethical Treatment of Non-Human Primates.

### Behavioral observations

Observations of sniffing behavior were conducted between October 17, 2020, and December 6, 2020, and thus, from the onset of the mating season to its peak (Fooden 2007). We observed 48 focal animals from two of the three groups. Focal animals comprised 24 females and 24 males from ages 1 to 25 (supplementary table S1), and thus covered all age classes, from juveniles (up to 2.5 years of age, N = 6) and subadults (up to 4.5 years of age, N = 9) to adults (5 years and older, N = 33, Kuester et al. 1995; Paul & Kuester 1990). 13 of 18 adult focal females had hormone implants. Each focal animal was observed six times for 20 minutes over the study period (total observation time = 48*2 hours = 96 hours). For each focal animal, three of six observations occurred in a feeding context (defined as at least ten minutes of feeding per protocol) and three in a non-feeding context. We spread the timing and context of observations per focal animal as equally as possible over the study period by observing the majority of individuals once before moving on to the next set of observations, and by conducting the subsequent observation of a given individual in a different context than the previous one whenever possible. During the focal protocols all instances of sniffing as well as details about the targets of sniffing were recorded. Observations were randomly distributed across the available daylight hours from 08:00 am to 05:00 pm. Because our study involved focal animals in the field, it was not possible to record data blind.

All focal observations were recorded with a digital video camera (Panasonic HC-V180) by one observer (M.S.). If the recorded subject moved out of sight during filming, the video was usually discarded. In 11 instances, two shorter videos were combined as one observation if the first video was of substantial length (around ten minutes). Seven individuals were juveniles who proved to be harder to film as they were more playful and tended to climb the trees out of sight, while three individuals were adult males that were rather shy and more difficult to observe. For these individuals we combined videos if the second video was recorded in a similar behavioral context, on the same day and was of sufficient length to complete the 20 min observation. For eight out of the 11 instances filming was resumed within 30 minutes, and for the remaining three instances up to 4.5h after the end of the first video. Sniffs were scored from the videos using the recorded video image as well as commentaries verbally recorded onto the video during the focal observations (see Tab. 1 for ethogram). We followed Jänig et al. (2018) in assigning each sniff recorded in the videos to one of three target categories: food, social (sniffs directed at a conspecific or its excretions) or other (sniffs directed at the environment, human-made objects and self-sniffs). We also noted the target object of each observed sniff and the behavior of the monkey after the sniff. All videos were analyzed by one observer (M.S.), however, five percent of the data were also analyzed by an additional observer trained by M.S. to check for inter-observer reliability. The audio-track of the video was unavailable to the second coder to avoid bias. By comparing the respective observed sniffs, the intra-class correlation coefficient (ICC) was calculated and revealed good reliability (ICC=0.86; Koo & Li 2016).

**Table 1.** Ethogram of observed behavior in this study.

| Behavior | Description |
| --- | --- |
| **Sniff** | individual bringing their nose $\leq 3\text{cm}$ towards an object/conspecific/themselves or touching an object with the hand and then bringing the hand towards the nose |
| **visual inspection** | males focusing on the female anogenital swelling from a close range $\leq 1\text{m}$ |
| **olfactory inspection** | a sniff (as defined above) directed at the female anogenital swelling |
| **copulation** | male mounting female and thrusting |

Besides the focal video protocols, *ad libitum* data were collected for every sniff that was observed outside the focal observations for focal and non-focal animals. Furthermore, whenever inspections of female swellings by males were observed, the following data were noted *ad libitum*: male and female ID, group, date, time, whether visual and/or olfactory inspection occurred and which inspection happened first, and if inspection was followed by a copulation (Tab. 1).

**Table 2.** Number of sniffs observed at different targets for 48 focal animals in 96 hours of focal observations.

| Sniffer sex | Food | Social | Other | Sum |
| --- | --- | --- | --- | --- |
| Male | 150 | 29 | 18 | **197** |
| Female | 274 | 11 | 29 | **314** |
| Sum | **423** | **40** | **47** | **511** |

### Statistical Analysis

Statistical analysis was conducted in R version 4.0.3 (R core team 2020) using Generalized Linear Mixed Models (GLMM) to allow accounting for repeated observations of the same individuals (Bates et al. 2015). We conducted three sets of models that were fitted by using the function ‘glmer’ from the package ‘lme4’ version 1.1-31 (Bates et al. 2015).

#### Model A: Sniffing frequencies

The aim of model A was to investigate the influence of sex, age and context on sniffing frequencies and, hence, to test our predictions 1), 2) and 3). Accordingly, we used ‘number of sniffs’ as the response variable, fitted with a Poisson error distribution. To be able to compare sniffing frequencies between our predictors, we extracted three sniffing frequencies, i.e., the number of sniffs at food, the number of sniffs at social targets and the number of sniffs at other targets separately from each 20min focal observation (N = 288 focal protocols x 3 target categories = 864 sniffing frequencies). Data collected *ad libitum* was not used in this model. Sex, age (years), observational context and target category (‘food’, ‘social’ or ‘other’) were fitted as fixed effects test predictors. Observational context referred to the context ‘feeding’ or ‘non-feeding’ in which each protocol was recorded, while target category referred to the object that was actually sniffed at, irrespective of whether the general observational context was feeding or not. Group, daytime and Julian day were fitted as fixed effects control predictors. Daytime was coded as morning (until 12:30 pm) or afternoon (after 12:30pm). Julian day was included to control for the progress of the mating season. Several two-way interactions were included in the model: I) sex and target category to address the prediction that males sniff more in a social setting and females more on food; II) sex and Julian day, since the observational period started at the early beginning of the mating season and ended at its height. Therefore, male and female sniffing behavior could show different patterns from the beginning to the end of the observation period; III) observational context and target category, because the observational contexts ‘feeding’ or ‘non-feeding’ presumably influence the probable targets of sniffing; IV) age and target category, to account for the possibility that age affects which objects the monkeys sniff at, and V) daytime and target category, as fresh food got distributed every morning at 08:00 and we accordingly expected more sniffs at edible items in the morning than in the afternoon. Individual ID and the ID of the observational protocol were included as a random effects control terms. To achieve more reliable p-values (Barr et al. 2013), we fitted random slopes of all predictors showing sufficient variation within the respective random intercept, i.e., the random slopes of Julian day, context*target category, and daytime*target category within individual ID, and target category within protocol ID.

#### Model B: Olfactory inspection of females

The aim of model B was to investigate the influence of female and male characteristics and the progression of the mating season on whether male inspections of female sexual swellings included olfaction or not (N = 176 genital inspections) and therefore our predictions 4) and 5). For this purpose, we fitted a GLMM with binominal error distribution using all genital inspections observed during focal observations as well as those recorded during *ad libitum* sampling, comprising data from 35 sexually mature and 3 immature males and from 37 sexually mature females. Female age (linear and squared) and male age, whether or not the female was contracepted and Julian day were fitted as fixed effects test predictors, while daytime and group were included as fixed effects control predictors. The two-way interactions of male age and female age (linear and squared), respectively, were included to account for the possibility that an effect of male age on olfactory inspection could be modulated by the age of the female partner in a linear or non-linear fashion. We also included an interaction between male age and contraception to address the possibility that inexperienced males might be more prone to gather additional cues about female fertility. Female and male ID were included as random effects control terms. For the random effect of male ID, the random slopes of Julian day, female age and contraception were incorporated. For female ID, the random slopes of Julian day and male age were included.

#### Model C: Genital inspections and copulation

Using focal and *ad libitum* data as described for model B, model C was fitted with binominal error distribution to investigate the influence of female and male characteristics and the occurrence of olfactory inspection on whether or not a genital inspection was followed by a copulation (N = 176 genital inspections) and, hence, test prediction 6. Female and male age, contraception (yes/no) and olfactory inspection (yes/no) were fitted as fixed effects test predictors while daytime, group and Julian day were included as fixed effects control predictors. We initially incorporated four two-way interactions: I) female and male age; II) female age and olfactory inspection; III) Julian day and olfactory inspection and IV) contraception and olfactory inspection. However, the interaction terms were too imbalanced, which caused stability issues (see general model procedures for stability checks), and were therefore removed again from the model. After removing all interactions, the model showed no stability issues. Female and male subject were included as random effects control terms. For the random effect of male ID, we included Julian day, olfactory inspection, contraception and female age as random slopes. For female ID, we included Julian day, olfactory inspection and male age as random slopes.

#### General Model Procedures

For all models, covariates were z-transformed to a mean of zero and a standard deviation of one before running the models to facilitate interpretation of model coefficients and model convergence (Schielzeth 2010). Variance Inflation Factors (VIF) were computed using the function ‘vif’ of the package ‘car’ (Fox & Weisberg 2019) to check for collinearity between the predictors (Quinn & Keough 2002). With largest VIFs of 1.08 (model A), 1.3 (model B) and 1.1 (model C), no collinearity issue could be detected in either model. All three models were tested for over- and underdispersion, with resulting dispersion parameters of 0.12 (model A), 0.87 (model B), and 0.65 (model C). As model A and C were underdispersed, the computed p-values should be considered as conservative (i.e. potentially biased towards higher p-values). Model stabilities were assessed by excluding levels of random effects one at a time. As mentioned above, with the exception of model C when fitted with interactions, none of the models showed stability issues.

A Likelihood Ratio Test (LRT) was used to determine the effects of the test predictors on the response variables by comparing the null model without fixed effects test predictors to the respective full model containing all predictors (Dobson, 2002; Forstmeier & Schielzeth 2011). If the full-null model comparison was significant (p < 0.05) or a trend (p < 0.1), the p-values of the individual predictors were afterwards determined by using the function ‘drop1’ from the package ‘lme4’ version 1.1-31. This function determines p-values by dropping single terms from the model and compares results with and without the respective predictor using LRTs (Bates et al. 2015). We computed pseudo-R^2^ values for Generalized Mixed-Effect models for the fixed effects (marginal R^2^) and for all model terms (conditional R^2^) to assess how much variance is explained by the models. R^2^ values were computed using the function ‘r.squaredGLMM’ from the package ‘MuMIn’ version 1.47.1, using the recommended trigamma method for Poisson, and delta method for binomial models. To facilitate interpretation of the main terms, interactions with a p > 0.1 were removed from the models, but only after conducting the respective full-null model comparison to ensure that predictions were tested with all predictors included. We report results for the dropped interactions in Tables 3 and 4. We computed 95% confidence intervals for all terms that remained in the model with parametric bootstrapping (n = 1000 simulations) using the ‘bootMer’ function from the package ‘lme4’ version 1.1-31.

**Table 3.** Results of the GLMM (model A) investigating sniffing frequency (number of sniffs) per focal animal, observation period and target category as response variable with Poisson error distribution (N = 288 observations x 3 target categories = 864). Terms in parentheses indicate trait levels relative to the respective reference level. Values for dropped interactions represent values before removal of the respective terms from the model. SE = standard error, CI = confidence interval. χ² and P values are derived from Likelihood Ratio Tests to determine the significance of the individual test predictors

| term | Estimate | SE | CI 2.5 | CI 97.5 | $\chi^2$ | P |
| --- | --- | --- | --- | --- | --- | --- |
| Intercept | -0.088 | 0.223 | -0.523 | 0.353 | † | † |
| **test predictors** |  |  |  |  |  |  |
| sex (male) | -0.588 | 0.207 | -1.035 | -0.182 | † | † |
| target category (other) | -6.941 | 1.469 | -8.477 | -6.43 | † | † |
| target category (social) | -6.177 | 1.188 | -8.43 | -6.13 | † | † |
| age | -0.374 | 0.108 | -0.549 | -0.173 | 10.991 | 0.001 |
| observ. context (non-feeding) | -1.160 | 0.210 | -1.491 | -0.689 | 31.154 | <0.001 |
| **control predictors** |  |  |  |  |  |  |
| group (2) | 0.105 | 0.205 | -0.258 | 0.462 | 0.265 | 0.606 |
| Julian day | -0.302 | 0.110 | -0.477 | -0.098 | 7.528 | 0.006 |
| daytime (morning) | 0.768 | 0.192 | 0.355 | 1.071 | 13.087 | <0.001 |
| **interactions** |  |  |  |  |  |  |
| target category*sex |  |  |  |  | 4.697 | 0.096 |
| target category (other)*sex (male) | 0.493 | 2.185 | -0.237 | 1.562 |  |  |
| target category (social)*sex (male) | 1.739 | 1.424 | 0.302 | 2.172 |  |  |
| **dropped interactions** |  |  |  |  |  |  |
| sex (male)*Julian day | -0.274 | 0.197 |  |  | 2.019 | 0.155 |
| observ. context*target category |  |  |  |  | 4.482 | 0.106 |
| context (non-feeding)*target (other) | 2.015 | 0.931 |  |  |  |  |
| context (non-feeding)*target (social) | 0.071 | 1.000 |  |  |  |  |
| age*target category |  |  |  |  | 0.707 | 0.702 |
| age*target (other) | -0.335 | 0.513 |  |  |  |  |
| age*target (social) | -0.228 | 0.430 |  |  |  |  |
| daytime*target category |  |  |  |  | 2.591 | 0.274 |
| daytime(morning)*target (other) | -1.141 | 0.892 |  |  |  |  |
| daytime(morning)*target (social) | -0.787 | 0.732 |  |  |  |  |

† not presented because of having a very limited interpretation

**Table 4.** Results of the binomial GLMM (model B) investigating the probability of an olfactory inspection during genital inspections (N = 176 inspections). Values in parentheses indicate trait levels relative to the respective reference level. Values for dropped interactions represent values before removal of the respective terms from the model. SE = standard error, CI = confidence interval. χ² and P values are derived from Likelihood Ratio Tests to determine the significance of the individual test predictors

| Term | Estimate | SE | CI 2.5 | CI 97.5 | $\chi^2$ | P |
| --- | --- | --- | --- | --- | --- | --- |
| Intercept | -1.7391 | 0.726 | -5.981 | -0.505 | † | † |
| **test predictors** |  |  |  |  |  |  |
| male age | -1.2686 | 0.5249 | -4.826 | -0.563 | † | † |
| female age | 0.425 | 0.2628 | -0.071 | 1.271 | 2.461 | 0.117 |
| female age<sup>2</sup> | 0.2542 | 0.1708 | -0.163 | 0.861 | 2.041 | 0.153 |
| contraception (yes) | 0.7468 | 0.7714 | -0.762 | 4.872 | † | † |
| **control predictors** |  |  |  |  |  |  |
| Julian day | -0.1512 | 0.2441 | -0.854 | 0.445 | 0.355 | 0.552 |
| daytime (morning) | 0.222 | 0.4129 | -0.757 | 1.181 | 0.271 | 0.603 |
| group (2) | -0.715 | 0.5745 | -2.208 | 0.677 | 1.269 | 0.260 |
| **interactions** |  |  |  |  |  |  |
| male age*contraception (yes) | 1.0138 | 0.599 | 0.01 | 4.366 | 2.998 | 0.083 |
| **dropped interactions** |  |  |  |  |  |  |
| male age*female age | 0.363 | 0.262 |  |  | 1.954 | 0.162 |
| male age*female age<sup>2</sup> | 0.018 | 0.197 |  |  | 0.009 | 0.925 |

† not presented because of having a very limited interpretation

## Results

In total, 511 sniffs were observed across all focal observations (96 hours total observation time), with 1.78 ± 1.70 (mean ± SD) sniffs per individual per 20 min observation period (corresponding to 5.34 sniffs per hour). Females were observed to sniff a total of 314 times (mean ± SD: 6.54 ± 8.7 per hour) and males 197 times (mean ± SD: 4.11 ± 5.43 per individual per hour, see Tab. 2). The vast majority of sniffs (83%) observed during focal observations was directed at food items (423/511), while 8% (40/511) and 9% (47/511) were directed at social or other targets, respectively (Tab. 2).

In the target context ‘food’, 49 different identifiable edible items (see supplementary table S2) were observed to be inspected via olfaction by the monkeys. In 288 (68%) of the 423 cases, the edible item was eaten after sniffing it, while it was discarded in the remainder. When food items had already been bitten into or torn apart (observable in 128 cases), they were eaten 89 times and thrown away 39 times after sniffing them. Items that appeared to be fresh (observable in 27 cases) were eaten 20 times and thrown away 7 times after sniffing. No corresponding data were collected on edible items that were not sniffed at, hence these results are only of a descriptive nature and were not further analyzed statistically.

Social sniffs were directed at a female swelling in 70% (28/40) of cases (26 by adult males, 2 by adult females). Eleven of the social sniffs were directed at an infant (1 by an adult male, 7 by adult females and 3 by another infant). One sniff was directed at a juvenile male by a five-year old male. Of the 47 sniffs at ‘other’ targets, 27 were self-sniffs, almost exclusively directed at their own hand after scratching themselves, 14 were directed at the environment (e.g. tree branch or ground) and 6 at human-made objects (e.g. bottle cap, camera trap).

In addition, 136 sniffs were observed *ad libitum*, of which 70 were directed at food, 44 at conspecifics and 22 at other targets. Of the 136 sniffs, 41 were observed for infants too young to identify. 87 sniffs were observed for identifiable individuals, with distributions across sex and age classes similar to data recorded during focal observations. Moreover, 52 sniffs were recorded for females, with 36 being directed at food, 8 at conspecifics, mostly infants, and 13 at themselves or the environment. Finally, 30 sniffs were recorded for males, with 16 being directed at food, 10 at conspecifics and four at themselves or the environment. Sniffs observed *ad libitum* included 18 cases of infants sniffing at their mother’s, or in one case their grandmother’s, mouth while they were eating.

### Model A: Sniffing frequencies

The test predictors in model A had a significant effect on the number of sniffs at a given target per individual per observation (LRT, N = 864, χ2 = 203.45, df = 14, P < 0.001, marginal R^2^ = 0.309, conditional R^2^ = 0.864). Specifically, sniffing frequencies differed between the sexes, whereby the effect of sex tended to depend on the target of the sniff. In particular, more sniffs were directed at food and ‘other’ targets by females compared to males. Males, on the other hand, sniffed more often at social targets than females (see Tab. 2 & 3, Fig. 1). We observed more sniffs during feeding than during non-feeding focal follows. Furthermore, younger individuals sniffed significantly more often than older individuals (Tab. 3, Fig. 2).

**Fig. 1.**
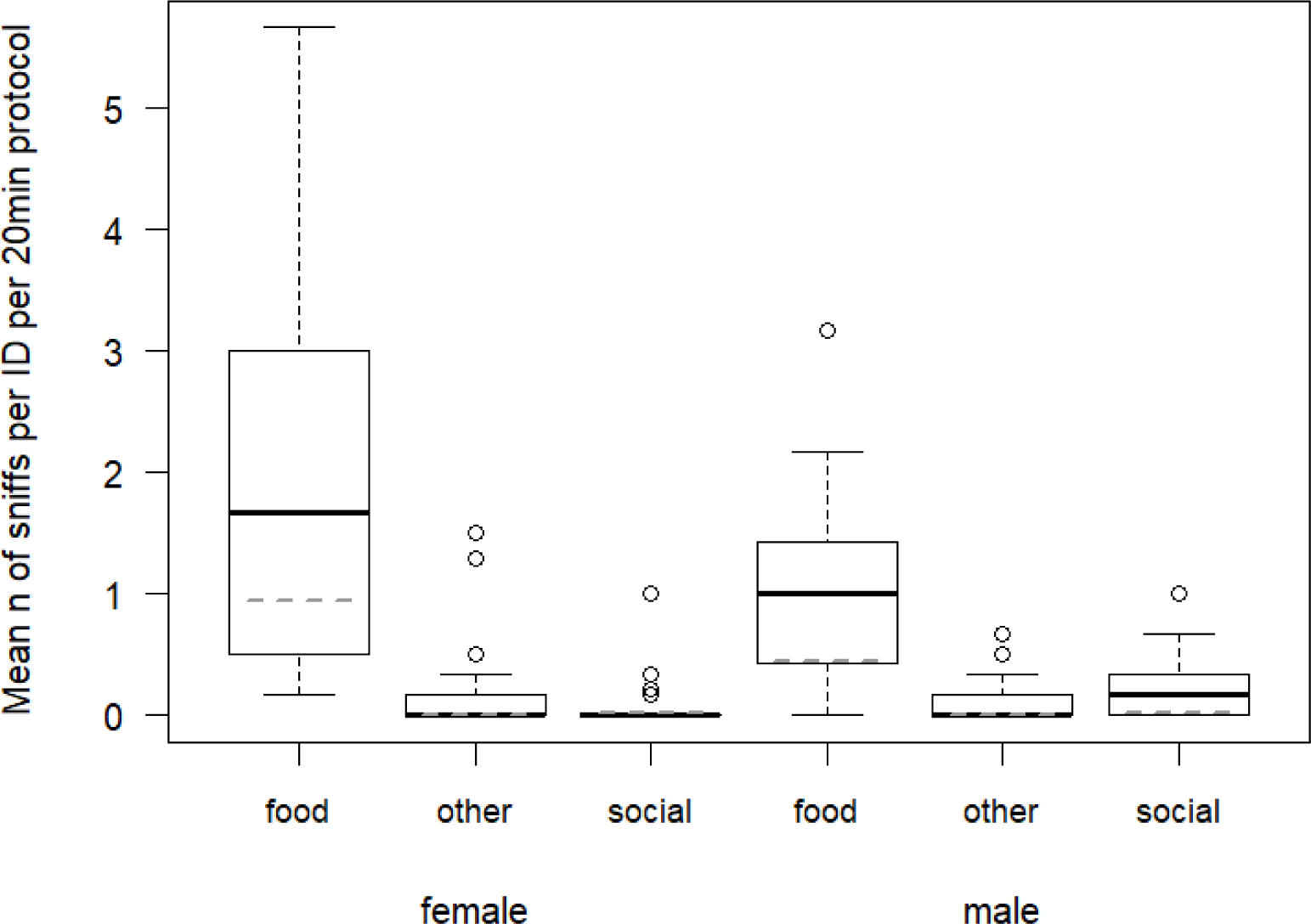
Mean number of sniffs per individual per 20 min focal observation for both sexes divided into the target categories food, other and social. Boxes represent medians and first and third quartiles, while the whiskers represent 1.5 times the interquartile range. Outliers are represented by data points. The horizontal dashed lines represent the model estimates when all other predictors are at their average.

**Fig. 2.**
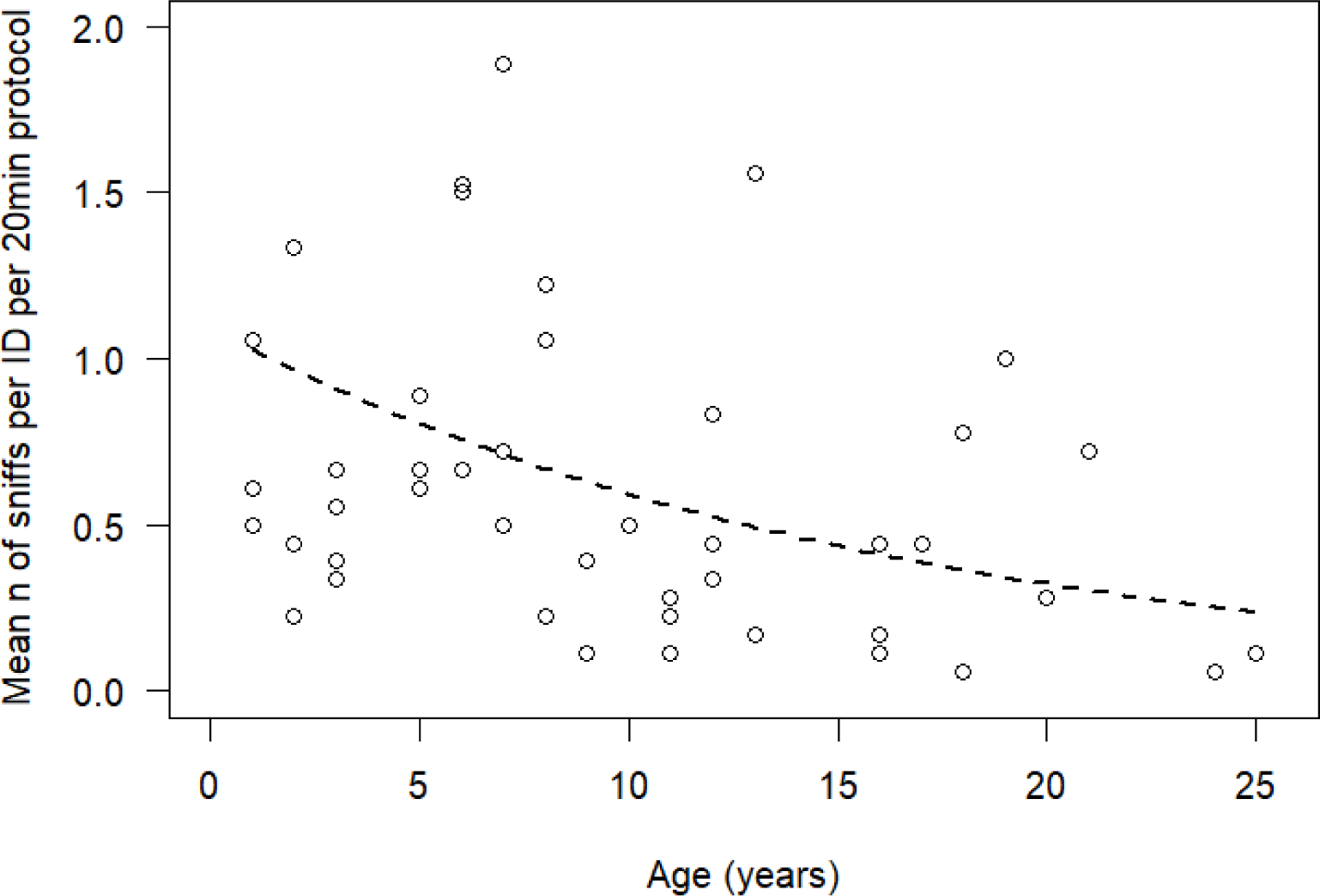
Mean number of sniffs per individual per 20 min focal observation across age. The dashed line represents the model estimate when all other predictors are at their average

Of the control predictors, Julian day and daytime significantly affected sniffing frequencies (see Tab. 3). In particular, we observed more sniffs earlier in the season and in the morning than in the afternoon. None of the other predictors had a significant influence on sniffing frequencies (Tab. 3).

### Model B: Olfactory inspection of females

Data on male inspections of sexual swellings, collected *ad libitum* as well as from focal observations, showed that out of 176 observed visual inspections, 51 were accompanied by an olfactory inspection of the swelling. The full-null comparison of model B investigating which parameters affected the occurrence of olfactory inspections revealed a trend (LRT, N = 176, χ2 =15.262, df = 8, P = 0.054, marginal R^2^ = 0.212, conditional R^2^ = 0.411). Specifically, the interaction between male age and contraception tended to influence the probability of an olfactory inspection (Tab. 4), whereby younger males generally tended to sniff slightly more during an inspection than older ones, but this effect was more pronounced when inspecting non-contracepted females (Fig. 3).

**Fig. 3.**
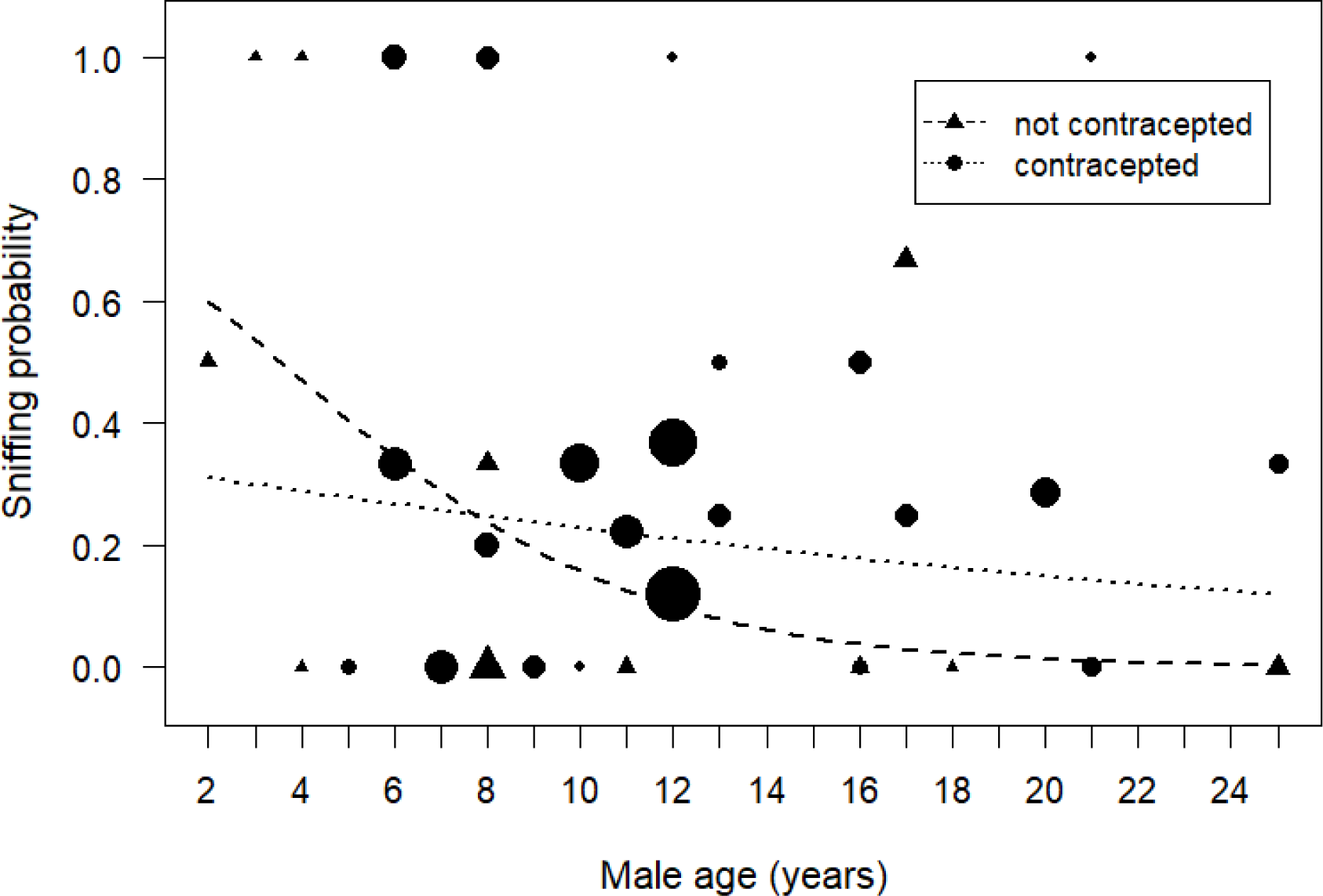
Sniffing probability during an anogenital inspection as a function of male age and female contraception (triangles: non-contracepted, circles: contracepted females). The lines represent the model estimate when all other predictors are at their average (dashed line: non-contracepted females, dotted line: contracepted females). Symbols are scaled relative to sample size (range 1-25 inspections).

### Model C: Genital inspections and copulation

Whether genital inspections were followed by a copulation was significantly affected by the suite of test predictors in model C (after excluding interactions from model due to stability issues (see methods), LRT, N = 176, χ2 = 31.951, df = 4, P < 0.001, marginal R^2^ = 0.357, conditional R^2^ = 0.513). In particular, olfactory inspection of the swelling significantly decreased the probability of copulation after inspection (Tab. 5). In fact, only 2 of the 51 visual *and* olfactory inspections were followed by a copulation, while copulation followed genital inspection in 44 of the 125 solely visual genital inspections. Furthermore, female age also influenced the copulation probability, with copulations more likely after inspections of younger females than older females (Tab. 5).

**Table 5.** Results of the binomial GLMM (model C) investigating the probability of a copulation following a genital inspection (N = 176 inspections). Values in parentheses indicate trait levels relative to the respective reference level. SE = standard error, CI = confidence interval. χ² and P values are derived from Likelihood Ratio Tests to determine the significance of the individual test predictors

| Term | Estimate | SE | CI 2.5 | CI 97.5 | $\chi^2$ | P |
| --- | --- | --- | --- | --- | --- | --- |
| Intercept | -0.280 | 0.705 | -2.452 | 1.701 | † | † |
| **test predictors** |  |  |  |  |  |  |
| olfactory inspection (yes) | -3.330 | 1.020 | -31.929 | -2.326 | 22.169 | <0.001 |
| male age | -0.104 | 0.287 | -1.061 | 0.671 | 0.120 | 0.729 |
| female age | -0.862 | 0.339 | -2.550 | -0.182 | 5.268 | 0.022 |
| contraception (yes) | -0.708 | 0.810 | -3.506 | 1.311 | 0.660 | 0.417 |
| **control predictors** |  |  |  |  |  |  |
| Julian day | -0.486 | 0.314 | -1.478 | 0.266 | 1.860 | 0.173 |
| daytime (morning) | -0.516 | 0.467 | -2.184 | 0.651 | 1.001 | 0.317 |
| group (2) | 0.411 | 0.645 | -0.968 | 2.558 | 0.321 | 0.571 |

† not presented because of having a very limited interpretation

## Discussion

This study contributes to understanding the sniffing behavior of Barbary macaques and thereby, catarrhine primates in general. In particular, we show that the frequency of sniffs varied with sex, age and context, in line with the hypothesis that individuals who benefit more from olfactory information than others sniff more frequently.

With an average of 5.3 sniffs per hour and individual, Barbary macaques sniffed at similar rates as some guenon species (*Cercopithecus diana, neglectus* and *hamlyni*) observed in captivity (6.1 sniffs/h, Zschoke & Thomsen 2014). In contrast, sniffing rates reported for chimpanzees in captivity were considerably lower (0.2 sniffs/h, Jänig et al. 2018) and even lower for chimpanzees observed in the wild (0.04 - 0.11 sniffs/h depending on season, Matsumoto-Oda et al. 2007). Furthermore, sniffing frequencies in Barbary macaques were not constant, but decreased over the course of the study. Along with different sampling conditions (wild *vs.* captivity, focal *vs.* group sampling) and the sparse availability of similar studies with different primate species, this makes meaningful comparisons between species difficult, calling for more studies on a wider range of species to assess interspecific patterns.

### Influence of context & target

In our study, the vast majority of sniffs were directed at food, which parallels findings for great apes (Jänig et al. 2018), guenons (Zschoke & Thomsen 2014) as well as mandrills and olive baboons (Laidre 2009). Primates have been suggested to steadily rely on olfactory cues to find and identify ripe food (e.g., Nevo & Heymann 2015), obtain information about nutritional value (Dominy et al. 2001), determine food safety and notice possible contamination (Sarabian et al. 2020). Hence, it comes as no surprise that an almost exclusively herbivorous species like the Barbary macaque shows frequent olfactory assessment of edible items. We were unable, however, to systematically assess why certain items were sniffed and others were not. Captive squirrel monkeys (*Saimiri sciureus)* and spider monkeys (*Ateles geoffroyi*) were described to rely mostly on visual cues when assessing familiar food items whereas novel food was more likely to prompt olfactory inspections (Laska et al. 2007b). In line with this suggestion, we observed a disproportional number of sniffs at hay and a chestnut provided by the park (data not shown), both of which were only rarely available to the monkeys. However, we did not systematically collect data on how often which types of food and other plants were available and therefore cannot address whether Barbary macaques really sniffed more frequently at novel or rare food items in this study.

### Influence of sex and age

The tendency that an effect of sex on sniffing probability is modulated by the target of a sniff supports prediction 1) that male Barbary macaques sniff more often in a social context than females do. Such an imbalance in social sniffs was also observed in other primate species such as chimpanzees (Jänig et al. 2018; Matsumota-Oda et al. 2007) and owl monkeys (*Aotus nancymaae*; Spence-Aizenberg 2017). Chimpanzees live in multi-male multi-female groups with fission-fusion dynamics and high levels of male-male competition for access to fertile females, enhancing the need for olfaction in the social context (Jänig et al. 2018; Matsumota-Oda et al. 2007). For Barbary macaques, which also live in multi-male multi-female groups, almost all social sniffs by adult males were observed to be directed at female sexual swellings and only one at another adult male. Thus, olfaction likely plays a role in male sexual competition for fertile females by, for example, providing information about the ovarian cycle, in addition to the visual signal of swelling size. However, sniffing does not seem as important for males to gather direct information about other males. Similarly, only in two instances a female was observed to sniff another female’s anogenital swelling. In both males and females, it could therefore be argued that an intra-sexual assessment of potential competitors either is not particularly relevant in such a highly promiscuous species, or that such an assessment is achieved using other sensory modalities. Possibly, aspects such as a male competitor’s size or female condition are more readily assessable using visual (Setchell et al. 2008; Tschoner 2015) or auditory cues (Pfefferle et al. 2008; Engelhardt et al. 2012).

Female Barbary macaques, on the other hand, sniffed more at food than males, supporting prediction 2). This corresponds to observations of female chimpanzees (Matsumota-Oda et al. 2007). Moreover, female primates of various species tend to be more wary than males of contamination risks for parasites or bacteria through food that is rotten or spoiled with feces (Sarabian et al. 2015; Sarabian et al. 2020; Poirotte et al. 2019), as evident from more olfactory inspection of contaminated food, followed by food manipulation (Sarabian et al. 2020). This behavior was rewarded by lower infection rates of females (Poulin 1996; Rolff 2002; Poirotte et al. 2019), suggesting that more visual or olfactory inspections, even though potentially costly, maximize the health, and therefore, also reproductive success of females. Similar mechanisms might apply to Barbary macaques, but a systematic analysis of parasite occurrence in both sexes would be needed to assess this possibility.

Furthermore, our prediction 3) of young individuals sniffing more often than older ones was supported with our data. These findings agree with the consensus that young animals inspect their environment more closely than older individuals because they are still in the process of learning to evaluate food, conspecifics or their general environment, and are comparable to observations in great apes (Jänig et al. 2018). Because the decrease of sniffing events with age in Barbary macaques appears to be quite linear rather than showing a sharp drop at old age (see Fig. 2), it is unlikely that it is primarily caused by a loss of olfactory capability at older age. Rather, this decrease may be caused by changes in experience and/or individual requirements. Particularly in the context of food assessment, sniffing frequencies may decrease as animals gain experience with different food items and conditions and learn to assess food quality based on visual or tactile cues. Additionally, it could be assumed that older females reaching post-reproductive age do not need to be as careful of contamination risks as females who still experience pregnancy and rear their young.

Together, 18 of 22 social sniffs that were observed for infant monkeys during *ad libitum* data collection were directed at the feeding mother’s, or in one case the grandmother’s, mouth. In each case the infants appeared to observe the eating behavior and tried to inspect the item visually. They were not chased off and the mothers reacted with indifference to the inspection attempt of their infant. This ‘muzzle-muzzle’ behavior has been observed in different mammals, including primates, and may enable the individuals to smell the breath of their conspecifics while they are eating (Nord et al. 2021; Arakawa et al. 2013; Laidre 2009). In this way they may gather information on which food has been deemed safe and valuable to eat by their conspecific.

### Olfaction in sexual interactions

Most social sniffs occurred in a sexual context and were directed at the female anogenital swelling, whereby about one third of inspections observed included sniffing. In line with prediction 4, younger males tended to sniff more frequently during an inspection than older ones, particularly if the inspected female was not contracepted. Although we observed contracepted females to show similar swelling sizes and patterns of tumescence and detumescence as non-contracepted ones (M.S., B.M.W. personal observations), this suggests that there may have been subtle differences in the visual appearance of the swellings that males could pick up upon. The nature of these differences and how they modulated sniffing behavior in an age-dependent fashion warrants further investigation. In contrast to prediction 5, however, we did not observe an effect of female age on the propensity of an olfactory inspection.

In line with prediction 6, the occurrence of an olfactory inspection affected subsequent sexual behavior. In particular, instances of sniffing the anogenital swelling almost exclusively led to no subsequent copulation. This may indicate that sniffing provides additional information about female fertile states or indicators of male competitors such as sperm remains that deters males from engaging in costly mating behavior, although male Barbary macaques were suggested not to be able to distinguish between fertile and non-fertile maximum swelling phases (Young et al. 2013). By inhibiting some males from inspecting their anogenital swelling close range, females may even manipulate the degree of access to olfactory information depending on the characteristics of the male.

Our assessment of olfactory inspections and their consequences (and thus of addressing prediction 4, 5 and 6) was certainly limited, because we did not track changes in female swelling size, ovarian hormone levels or whether observed copulations actually led to ejaculation. Additionally, female odor cues could potentially be gathered by males from a greater distance than what we defined as a sniff in this study. It is possible that the active sniffs we scored were associated with situations in which olfactory cues were so subtle that they required bringing the nose close to the odor source, while at other times, olfactory cues may have been strong enough to be perceivable from a greater distance, requiring no movement towards the odor source that is visible to the observer. We therefore are not able to entirely rule out that what we scored as visual inspections were truly multimodal ones that included a potential olfactory component. Whether this could be the case is not assessable from observations alone, and would require an experimental approach. As we only ever observed an active protruding out of the nose from a very close distance to an object and not from further away, however, we believe that our definition covers the vast majority of instances of active olfactory sampling.

Unfortunately, the low number of olfactory inspections that led to copulation did not allow us to address whether aspects such as female age or hormonal contraception modulated an effect of olfactory inspection on copulation probabilities. Studies on other primate species report differences in mating behavior or olfactory signals related to hormonal contraception (Japanese macaques, *M. fuscata*, Leca et al. 2018, ring-tailed lemurs *Lemur catta*, Crawford et al. 2011). By contrast, owl monkeys (*Aotus nancymaae*) showed no change in the released chemical profiles of females with contraception and no difference in the ability of females to form new pair bonds with males (Spence-Aizenberg et al. 2018). Our results provide no indication that female contraception affected the likelihood of a copulation after inspection, but the limitations indicated above prevent us from drawing more detailed conclusions about the role of sniffing in Barbary macaque mating interactions, which would require more comprehensive focal observations of sexual interactions, including an assessment of female fertility based on hormonal data. A greater focus on sniffing in a social context along with data on dominance and other social interactions further would open up possibilities to address the role of olfaction in intra-sexual competition.

In conclusion, Barbary macaques routinely sniff in different contexts and sniffing behavior was modulated by individual attributes such as sex and age. These findings are in line with current research in other (catarrhine) primates and add to the growing evidence about the importance of olfaction across primate species. Subsequent research is needed to thoroughly interpret sniffing behavior at food or conspecifics in the light of visual or other available sensory information, with this study serving as a basis. As such, this study represents a first step towards understanding the use and importance of olfaction in the social lives of Barbary macaques.

## Acknowledgements

This study was funded by the German Research Foundation, DFG (grant no. SCHL 2011/2-1 awarded to B.M. Weiß).

We are highly grateful to E. Merz, R. & M. Hilgartner and the team at Affenberg Salem for permission to conduct the study, provisioning of demographic information and their continuing support throughout the study. We are also thankful to A. Krüger for coding videos for the assessment of inter-observer reliability and to the University of Leipzig for logistic support. R. Mundry provided R functions for assessing model assumptions and stability. Three reviewers provided valuable comments on an earlier version of the manuscript.

---

**Table S1:**
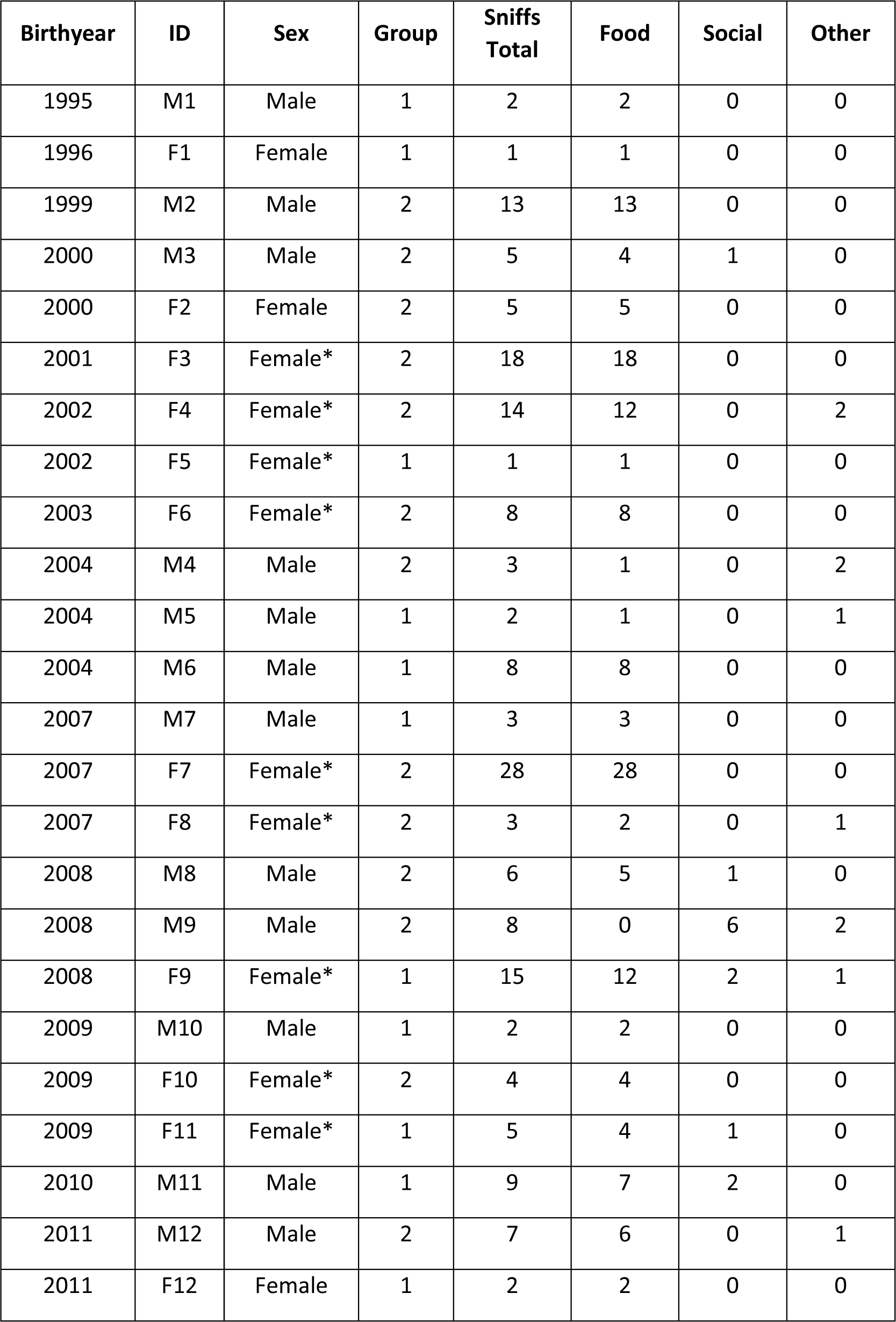
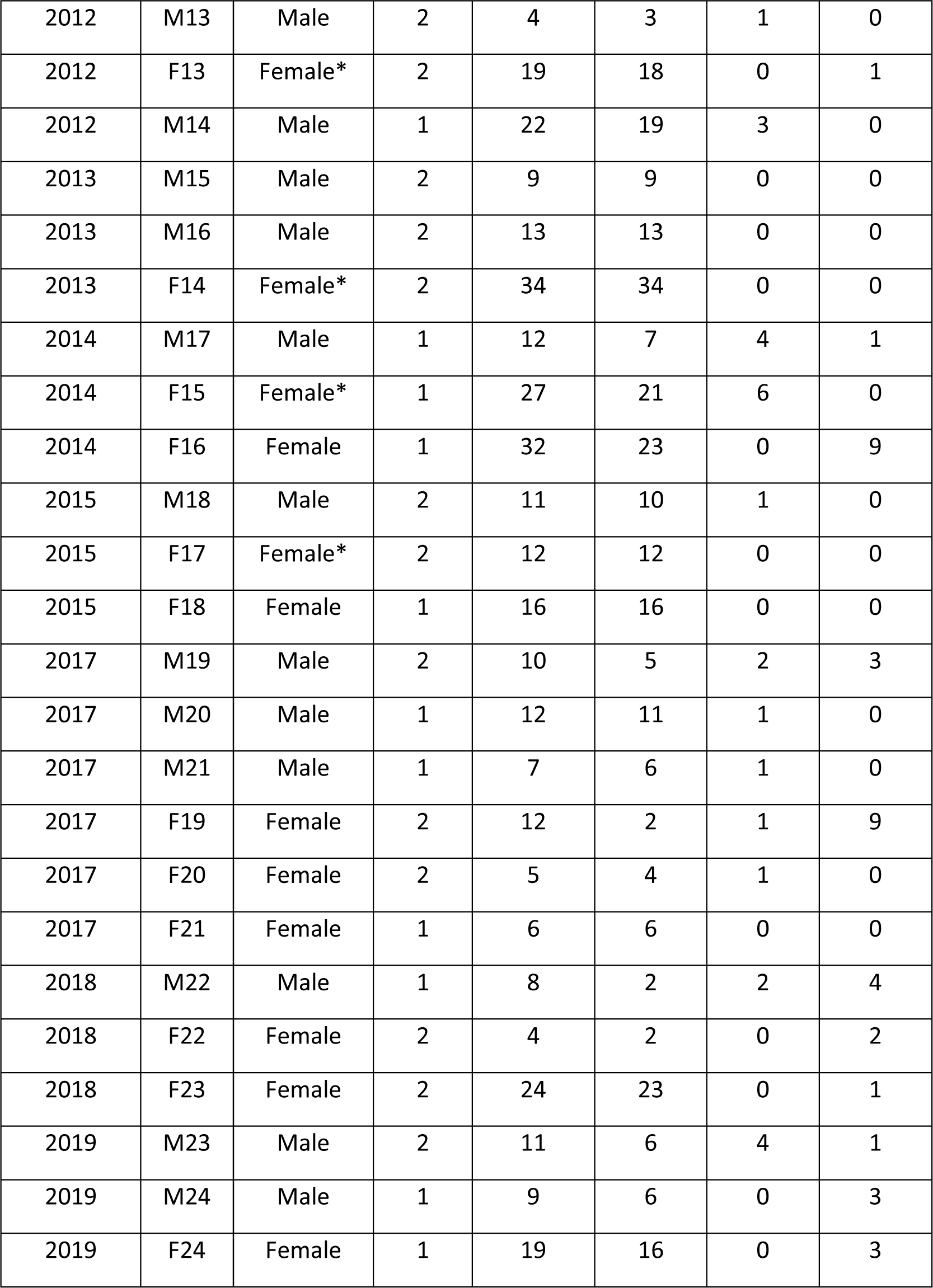
Overview of observed sniffs for 48 focal animals, including their birth year, sex, group, total sniffs and distribution of sniffs in feeding, social or other contexts. Asterisk notes female on contraception.

**Table S2:** Identified edible items focal animals sniffed at and the respective number of observed sniffs. Not included were unidentifiable objects.

| Item | No. Sniffs |
| --- | --- |
| Potato | 30 |
| Leaf | 26 |
| Carrot | 23 |
| Pineapple | 22 |
| Apple | 21 |
| Salad | 17 |
| Tomato | 16 |
| Leek | 15 |
| Grape | 14 |
| Hay | 11 |
| Popcorn | 11 |
| Parsley | 10 |
| Capsicum | 8 |
| Green onion | 8 |
| Beech nut | 7 |
| Chestnut | 7 |
| Stems | 7 |
| Celery | 6 |
| Orange | 6 |
| Seed | 6 |
| Tree branch | 6 |
| Mushroom | 5 |
| Broccoli | 4 |
| Cucumber | 4 |
| Fennel | 4 |
| Mango | 4 |
| Basil | 3 |
| Brussel sprout | 3 |
| Cabbage | 3 |
| Chinese cabbage | 3 |
| Honey melon | 3 |
| Potato "Vitelotte" | 3 |
| Banana | 2 |
| Black radish | 2 |
| Pear | 2 |
| Pomegranate | 2 |
| Walnut | 2 |
| Wheat | 2 |
| Avocado | 1 |
| Beetroot | 1 |
| Chives | 1 |
| Clover | 1 |
| Eggplant | 1 |
| Fern | 1 |
| Grass | 1 |
| Pellet ("monkey chow") | 1 |
| Red cabbage | 1 |
| Thyme | 1 |

## References

Arakawa, H., Kelliher, K. R., Zufall, F., & Munger, S. D. (2013). The receptor guanylyl cyclase type d (Gc-d) ligand uroguanylin promotes the acquisition of food preferences in mice. Chemical Senses, 38(5), 391–397. 10.1093/chemse/bjt015

Barr, D. J., Levy, R., Scheepers, C., & Tily, H. J. (2013). Random effects structure for confirmatory hypothesis testing: Keep it maximal. Journal of Memory and Language, 68(3), 255–278. 10.1016/j.jml.2012.11.001

Barton, R. A. (2006). Olfactory evolution and behavioral ecology in primates. American Journal of Primatology, 68(6), 545–558. 10.1002/ajp.20251

Bates, D., Mächler, M., Bolker, B., & Walker, S. (2015). Fitting linear mixed-effects models using lme4. Journal of Statistical Software, 67(1). 10.18637/jss.v067.i01

Boulet, M., Crawford, J. C., Charpentier, M. J., & Drea, C. M. (2010). Honest olfactory ornamentation in a female-dominant primate. Journal of Evolutionary Biology, 23(7), 1558–1563. 10.1111/j.1420-9101.2010.02007.x

Brauch, K., Pfefferle, D., Hodges, K., Möhle, U., Fischer, J., & Heistermann, M. (2007). Female sexual behavior and sexual swelling size as potential cues for males to discern the female fertile phase in free-ranging Barbary macaques (*Macaca sylvanus*) of Gibraltar. Hormones and Behavior, 52(3), 375–383. 10.1016/j.yhbeh.2007.06.001

Campos, F. A., Altmann, J., Cords, M., Fedigan, L. M., Lawler, R., Lonsdorf, E. V., Stoinski, T. S., Strier, K. B., Bronikowski, A. M., Pusey, A. E., & Alberts, S. C. (2022). Female reproductive aging in seven primate species: Patterns and consequences. Proceedings of the National Academy of Sciences of the United States of America, 119(20), e2117669119. 10.1073/pnas.2117669119

Crawford, J. C., Boulet, M., & Drea, C. M. (2011). Smelling wrong: hormonal contraception in lemurs alters critical female odour cues. Proceedings of the Royal Society B: Biological Sciences, 278(1702), 122–130. 10.1098/rspb.2010.1203

Cunningham, E. P., Edmonds, D., Stalter, L., & Janal, M. N. (2021). Ring-tailed lemurs (*Lemur catta*) use olfaction to locate distant fruit. American Journal of Physical Anthropology, 175(1), 300–307. 10.1002/ajpa.24255

de Turckheim, G., & Merz, E. (1984). Breeding Barbary macaques in outdoor open enclosures. In J. E. Fa (Ed.), The Barbary macaque: A case study in conservation (pp. 241–261). Plenum Publishing Corporation. 10.1007/978-1-4613-2785-1_10

Dobson, A. J. (2002). An introduction to generalized linear models (2nd ed). Chapman & Hall/CRC.

Dominy, N. J., Lucas, P. W., Osorio, D., & Yamashita, N. (2001). The sensory ecology of primate food perception. Evolutionary Anthropology: Issues, News, and Reviews, 10(5), 171–186. 10.1002/evan.1031

Engelhardt, A., Fischer, J., Neumann, C., Pfeifer, J. B., & Heistermann, M. (2012). Information content of female copulation calls in wild long-tailed macaques (*Macaca fascicularis*). Behavioral ecology and sociobiology, 66(1), 121–134. 10.1007/s00265-011-1260-9

Fooden, J. (2007). Systematic review of the Barbary macaque, *Macaca sylvanus* (Linnaeus, 1758). Fieldiana Zoology, 113(1), 1. 10.3158/0015-0754(2007)113[1:SROTBM]2.0.CO;2

Forstmeier, W., & Schielzeth, H. (2011). Cryptic multiple hypotheses testing in linear models: Overestimated effect sizes and the winner’s curse. Behavioral Ecology and Sociobiology, 65(1), 47–55. 10.1007/s00265-010-1038-5

Fox, J. & Weisberg, S. (2019). An {R} Companion to Applied Regression, Third Edition. Thousand Oaks CA: Sage. URL: https://socialsciences.mcmaster.ca/jfox/Books/Companion/

Henkel, S., Lambides, A. R., Berger, A., Thomsen, R., & Widdig, A. (2015). Rhesus macaques (*Macaca mulatta*) recognize group membership via olfactory cues alone. Behavioral Ecology and Sociobiology, 69(12), 2019–2034. 10.1007/s00265-015-2013-y

Henkel, S., & Setchell, J. M. (2018). Group and kin recognition via olfactory cues in chimpanzees (*Pan troglodytes)*. Proceedings of the Royal Society B: Biological Sciences, 285(1889), 20181527. 10.1098/rspb.2018.1527

Hiramatsu, C., Melin, A. D., Aureli, F., Schaffner, C. M., Vorobyev, M., & Kawamura, S. (2009). Interplay of olfaction and vision in fruit foraging of spider monkeys. Animal Behaviour, 77(6), 1421–1426. 10.1016/j.anbehav.2009.02.012

Janda, E. D., Perry, K. L., Hankinson, E., Walker, D., & Vaglio, S. (2019). Sex differences in scent-marking in captive red-ruffed lemurs. American Journal of Primatology, 81(1), e22951. 10.1002/ajp.22951

Jänig, S., Weiß, B. M., & Widdig, A. (2018). Comparing the sniffing behavior of great apes. American Journal of Primatology, 80(6), e22872. 10.1002/ajp.22872

Kay, R. F. (2018). 100 years of primate paleontology. American Journal of Physical Anthropology, 165(4), 652–676. 10.1002/ajpa.23429

Koo, T. K., & Li, M. Y. (2016). A Guideline of Selecting and Reporting Intraclass Correlation Coefficients for Reliability Research. Journal of Chiropractic Medicine, 15(2), 155–163. 10.1016/j.jcm.2016.02.012

Kücklich, M., Weiß, B. M., Birkemeyer, C., Einspanier, A., & Widdig, A. (2019). Chemical cues of female fertility states in a non-human primate. Scientific Reports, 9(1), 13716. 10.1038/s41598-019-50063-w

Kuester, J., & Paul, A. (1992). Influence of male competition and female mate choice on Male mating success in Barbary macaques (*Macaca sylvanus*). Behaviour, 120(3–4), 192–216. 10.1163/156853992X00606

Kuester, J., Paul, A. & Arnemann, J. (1995). Age-related and individual differences of reproductive success in male and female barbary macaques (*Macaca sylvanus)*. Primates, 36, 461–476. 10.1007/BF02382869

Laidre, M. E. (2009). Informative breath: Olfactory cues sought during social foraging among old world monkeys (*Mandrillus sphinx, M. Leucophaeus*, and *Papio anubis*). Journal of Comparative Psychology, 123(1), 34–44. 10.1037/a0013129

Laska, M., Bautista, R. M. R., Höfelmann, D., Sterlemann, V., & Salazar, L. T. H. (2007a). Olfactory sensitivity for putrefaction-associated thiols and indols in three species of non-human primate. Journal of Experimental Biology, 210(23), 4169–4178. 10.1242/jeb.012237

Laska, M., Freist, P., & Krause, S. (2007b). Which senses play a role in nonhuman primate food selection? A comparison between squirrel monkeys and spider monkeys. American Journal of Primatology, 69(3), 282–294. 10.1002/ajp.20345

Leca, J. B., Gunst, N., Shimizu, K., Huffman, M. A., Takahata, Y., & Vasey, P. L. (2018). Hormonal contraceptive affects heterosexual but not homosexual behavior in free-ranging female Japanese macaques over 17 mating seasons. Hormones and Behavior, 105, 166–176. 10.1016/j.yhbeh.2018.08.009

Matsui, A., Go, Y., & Niimura, Y. (2010). Degeneration of olfactory receptor gene repertories in primates: no direct link to full trichromatic vision. Molecular biology and evolution, 27(5), 1192–1200. 10.1093/molbev/msq003

Matsumoto-Oda, A., Kutsukake, N., Hosaka, K., & Matsusaka, T. (2007). Sniffing behaviors in Mahale chimpanzees. Primates, 48(1), 81–85. 10.1007/s10329-006-0006-1

Melin, A.D., Nevo, O., Shirasu, M., Williamson, R. E., Garrett, E. C., Endo, M., Sakurai, K., Matsushita, Y., Touhara, K., & Kawamura, S. (2019). Fruit scent and observer colour vision shape food-selection strategies in wild capuchin monkeys. Nature Communications, 10, 2407. 10.1038/s41467-019-10250-9

Modolo, L., & Martin, R. D. (2008). Reproductive success in relation to dominance rank in the absence of prime-age males in Barbary macaques. American Journal of Primatology, 70(1), 26–34. 10.1002/ajp.20452

Nevo, O., & Heymann, E. W. (2015). Led by the nose: Olfaction in primate feeding ecology. Evolutionary Anthropology: Issues, News, and Reviews, 24(4), 137–148. 10.1002/evan.21458

Niimura, Y., Matsui, A., & Touhara, K. (2018). Acceleration of olfactory receptor gene loss in primate evolution: Possible link to anatomical change in sensory systems and dietary Transition. Molecular Biology and Evolution, 35(6), 1437–1450. 10.1093/molbev/msy042

Nord, C. M., Bonnell, T. R., Dostie, M. J., Henzi, S. P., & Barrett, L. (2021). Tolerance of muzzle contact underpins the acquisition of foraging information in vervet monkeys (*Chlorocebus pygerythrus*). Journal of Comparative Psychology, 135(3), 349–359. 10.1037/com0000258

Paul, A., & Kuester, J. (1990). Adaptive significance of sex ratio adjustment in semifree-ranging Barbary macaques (*Macaca sylvanus*) at Salem. Behavioral Ecology and Sociobiology, 27(4), 287–293.

Paul, A., Kuester, J. & Podzuweit, D. (1993). Reproductive senescence and terminal investment in female Barbary macaques (*Macaca sylvanus*) at Salem. International Journal of Primatology, 14, 105–124. 10.1007/BF02196506

Petrulis, A., Peng, M., & Johnston, R. E. (1999). Effects of vomeronasal organ removal on individual odor discrimination, sex-odor preference, and scent marking by female hamsters. Physiology and Behavior, 66(1), 73–83. 10.1016/S0031-9384(98)00259-5

Pfefferle, D., Heistermann, M., Hodges, K. & Fischer, J. (2008). Male Barbary macaques eavesdrop on mating outcome: a playback study. Animal Behaviour. 75. 1885–1891. 10.1016/j.anbehav.2007.12.003.

Poirotte, C., Massol, F., Herbert, A., Willaume, E., Bomo, P. M., Kappeler, P. M., & Charpentier, M. J. E. (2017). Mandrills use olfaction to socially avoid parasitized conspecifics. Science Advances, 3(4), e1601721. 10.1126/sciadv.1601721

Poirotte, C., Sarabian, C., Ngoubangoye, B., MacIntosh, A. J. J., & Charpentier, M. (2019). Faecal avoidance differs between the sexes but not with nematode infection risk in mandrills. Animal Behaviour, 149, 97–106. 10.1016/j.anbehav.2019.01.013

Poulin, R. (1996). Sexual inequalities in helminth infections: A cost of being a male? The American Naturalist, 147(2), 287–295. 10.1086/285851

Quinn, G., & Keough, M. (2002). Experimental Design and Data Analysis for Biologists. Cambridge: Cambridge University Press. doi:10.1017/CBO9780511806384

R Core Team 2020. R: A language and environment for statistical computing. R Foundation for Statistical Computing, Vienna, Austria. www.R-project.org

Rigaill, L., Higham, J. P., Lee, P. C., Blin, A., & Garcia, C. (2013). Multimodal sexual signaling and mating behavior in Olive baboons (*Papio anubis*). American Journal of Primatology, 75(7), 774–787. 10.1002/ajp.22154

Rigaill, L., Vaglio, S., Setchell, J. M., Suda-Hashimoto, N., Furuichi, T., & Garcia, C. (2022). Chemical cues of identity and reproductive status in Japanese macaques. American Journal of Primatology, 84(8), e23411. 10.1002/ajp.23411

Rolff, J. (2002). Bateman’s principle and immunity. Proceedings of the Royal Society B: Biological Sciences, 269(1493), 867–872. 10.1098/rspb.2002.1959

Rushmore, J., Leonhardt, S. D., & Drea, C. M. (2012). Sight or scent: lemur sensory reliance in detecting food quality varies with feeding ecology. PloS One, 7(8), e41558. 10.1371/journal.pone.0041558

Sarabian, C., & MacIntosh, A. J. J. (2015). Hygienic tendencies correlate with low geohelminth infection in free-ranging macaques. Biology Letters, 11(11), 20150757. 10.1098/rsbl.2015.0757

Sarabian, C., Ngoubangoye, B., & MacIntosh, A. J. J. (2020). Divergent strategies in faeces avoidance between two cercopithecoid primates. Royal Society Open Science, 7(3), 191861. 10.1098/rsos.191861

Schielzeth, H. (2010). Simple means to improve the interpretability of regression coefficients: Interpretation of regression coefficients. Methods in Ecology and Evolution, 1(2), 103–113. 10.1111/j.2041-210X.2010.00012.x

Scordato, E. S., & Drea, C. M. (2007). Scents and sensibility: information content of olfactory signals in the ringtailed lemur, *Lemur catta*. Animal Behaviour, 73(2), 301–314. doi:10.1016/j.anbehav.2006.08.006

Setchell, J. M., Smith, T., Wickings, E. J., & Knapp, L. A. (2008). Social correlates of testosterone and ornamentation in male mandrills. Hormones and behavior, 54(3), 365–372. 10.1016/j.yhbeh.2008.05.00

Setchell, J. M., Vaglio, S., Moggi-Cecchi, J., Boscaro, F., Calamai, L., & Knapp, L. A. (2010). Chemical composition of scent-gland secretions in an old world monkey (*Mandrillus sphinx*): influence of sex, male status, and individual identity. Chemical senses, 35(3), 205–220. 10.1093/chemse/bjp105

Sipos, M. L., Wysocki, C. J., Nyby, J. G., Wysocki, L., & Nemura, T. A. (1995). An ephemeral pheromone of female house mice: Perception via the main and accessory olfactory systems. Physiology and Behavior, 58(3), 529–534. 10.1016/0031-9384(95)00089-2

Smith, T. D., & Bhatnagar, K. P. (2004). Microsmatic primates: Reconsidering how and when size matters. The Anatomical Record, 279B(1), 24–31. 10.1002/ar.b.20026

Spence-Aizenberg, A. M. (2017). Olfactory communication, mate choice, and reproduction in a pair bonded primate (*Aotus spp*.). Publicly Accessible Penn Dissertations. 2885.

Spence-Aizenberg, A., Kimball, B.A., Williams, L.E., & Fernandez-Duque, E. (2018). Chemical composition of glandular secretions from a pair-living monogamous primate: Sex, age, and gland differences in captive and wild owl monkeys (*Aotus spp*.). American Journal of Primatology, 80:e22730. 10.1002/ajp.22730

Suendermann, D., Scheumann, M., & Zimmermann, E. (2008). Olfactory predator recognition in predator-naïve gray mouse lemurs (*Microcebus murinus*). Journal of Comparative Psychology, 122(2), 146–155. 10.1037/0735-7036.122.2.146

Tschoner, S.E. (2015). Sexual swelling as honest signal for fertility in female Barbary macaques (Macaca sylvanus)? (Master Thesis) Universität Wien. Fakultät für Lebenswissenschaften.

Vaglio, S., Minicozzi, P., Kessler, S. E., Walker, D., & Setchell, J. M. (2021). Olfactory signals and fertility in olive baboons. Scientific Reports, 11(1), 8506. 10.1038/s41598-021-87893-6

Young, C., Majolo, B., Heistermann, M., Schülke, O., & Ostner, J. (2013). Male mating behaviour in relation to female sexual swellings, socio-sexual behaviour and hormonal changes in wild Barbary macaques. Hormones and Behavior, 63(1), 32–39. 10.1016/j.yhbeh.2012.11.004

Zschoke, A., & Thomsen, R. (2014). Sniffing behaviours in guenons. Folia Primatologica, 85(4), 244–251. 10.1159/000363409

